# Ribozyme-mediated gene-fragment complementation for non-destructive reporting of DNA transfer within soil

**DOI:** 10.1101/2024.04.05.588345

**Authors:** Malyn A. Selinidis, Andrew C. Corliss, James Chappell, Jonathan J. Silberg

## Abstract

Enzymes that produce volatile metabolites can be coded into genetic circuits to report non-disruptively on microbial behaviors in hard-to-image soils. However, these enzyme reporters remain challenging to apply in gene transfer studies due to leaky off states that can lead to false positives. To overcome this problem, we designed a reporter that uses ribozyme-mediated gene-fragment complementation of a methyl halide transferase (MHT) to regulate the synthesis of methyl halides. We split the *mht* gene into two non-functional fragments and attached these to a pair of splicing ribozyme fragments. While the individual *mht*-ribozyme fragments did not produce methyl halides when transcribed alone in *Escherichia coli*, co-expression resulted in a spliced transcript that translated the MHT reporter. When cells containing one *mht*-ribozyme fragment transcribed from a mobile plasmid were mixed with cells that transcribed the second *mht*-ribozyme fragment, methyl halides were only detected following rare conjugation events. When conjugation was performed in soil, it led to a 16-fold increase in methyl halides in the soil headspace. These findings show how ribozyme-mediated gene-fragment complementation can achieve tight control of protein reporter production, a level of control that will be critical for monitoring the effects of soil conditions on gene transfer and the fidelity of biocontainment measures developed for environmental applications.

## Introduction

Microorganisms evolve and adapt to their environment by taking up mobile DNA through conjugation, transduction, transformation, and vesicle fusion.^1,2^ These processes, described collectively as horizontal gene transfer (HGT), support microbial adaptation in dynamic environments by helping them acquire beneficial traits, like increased heat resistance, expanded catabolic capabilities, and enhanced resistance to toxins.^3–6^ HGT is also influenced by human activity, which has stimulated the spread of antibiotic-resistance.^1,4^ For example, the widespread use of antibiotics in farm animals has led to increased antibiotic-resistant microbes in soils,^7^ due to the exchange of antibiotic-resistance genes in these environments.^8^ In addition, HGT represents a risk for applications of synthetic biology in soil, such as biotechnologies being developed to accelerate bioremediation, fix nitrogen for agricultural applications, and store carbon to mitigate greenhouse gas emissions.^9–11^ The spread of synthetic genes in native microbial communities has the potential to result in phenotypes that alter consortia composition and the collective behaviors of the organisms in those communities. Despite the importance of understanding HGT in the environment and employing land use practices that minimize risks, our tools for monitoring HGT *in situ* remain limited.

Numerous techniques have been described for monitoring gene transfer,^3^ which can be separated into two major technology types. First, chemical methods have been developed, such as high-throughput chromosome capture (Hi-C) and Emulsion, Paired Isolation and Concatenation PCR (EpicPCR),^12,13^ which cross-link mobile and host DNA chemically and then use next-generation sequencing to identify who acquired the mobile DNA. Second, synthetic genetic circuits have been created that report on HGT by producing unique cellular outputs upon mobile DNA uptake. These outputs produce phenotypes, such as antibiotic resistance or fluorescence, which can be quantified using selections.^14,15^ Alternatively, synthetic genetic circuits can code information in the genome or transcriptome through DNA or RNA editing tools, which can be read out without selection using sequencing.^16,17^ When studying HGT in soil materials, these techniques all require cell or nucleic acid extraction prior to measurement, which can vary in efficiency with soil properties.^18^ As such, these approaches are limited in their ability to provide dynamic gene transfer information without disrupting the sample.

Enzymes that synthesize rare volatile gases can non-disruptively report on soil microbial behaviors. One such reporter is Methyl Halide Transferase (MHT), a rare enzyme that uses halide ions and S-adenosylmethionine as substrates to synthesize volatile methyl halides.^19^ These enzymes are appealing to use as reporters for soil synthetic biology because their volatile products diffuse out of soil matrices into the headspace where they can be quantified using gas chromatography.^20^ MHT reporters have been used for in soils to monitor the bioavailability of microbe-microbe signals^21^, microbe-plant signals^22^, and sugars.^23^ In addition, they have been used to monitor the conditional activities of soil microbe promoters^24^ and coded into mobile DNA to report on conjugation in soil.^25^ Unfortunately, MHT reporters remain limited in their ability to report on HGT in soils, because a background signal accumulates with time in the absence of HGT due to leaky MHT expression.

Here, we investigate whether stringent control over a gene transfer reporter can be achieved by implementing multi-level synthetic gene controls. A novel tool for controlling reporter protein production at the RNA level is the self-splicing ribozyme from *Tetrahymena thermophila*, which joins two RNAs found on either side of the ribozyme to form a spliced RNA output. Recently, an RNA engineering study identified sites where the ribozyme can be split into two fragments that are inactive unless combined using additional RNA interaction sequences.^26^ By combining the split ribozyme design with a split protein sequence (Figures 1A-C), we use ribozyme-mediated gene-fragment complementation to form a single RNA transcript and ultimately a single protein product through control at both the RNA and protein levels. By applying this approach to an MHT gas reporter, we show that this multi-level control abolishes reporter leak and allows for conjugation studies by creating a signal only when both *mht-*ribozyme fragments are present in a cell that participates in gene transfer. We show that this new methodology allows for exquisite control of gas production and is compatible with studies of gene transfer in complex soil materials.

**Figure 1.**
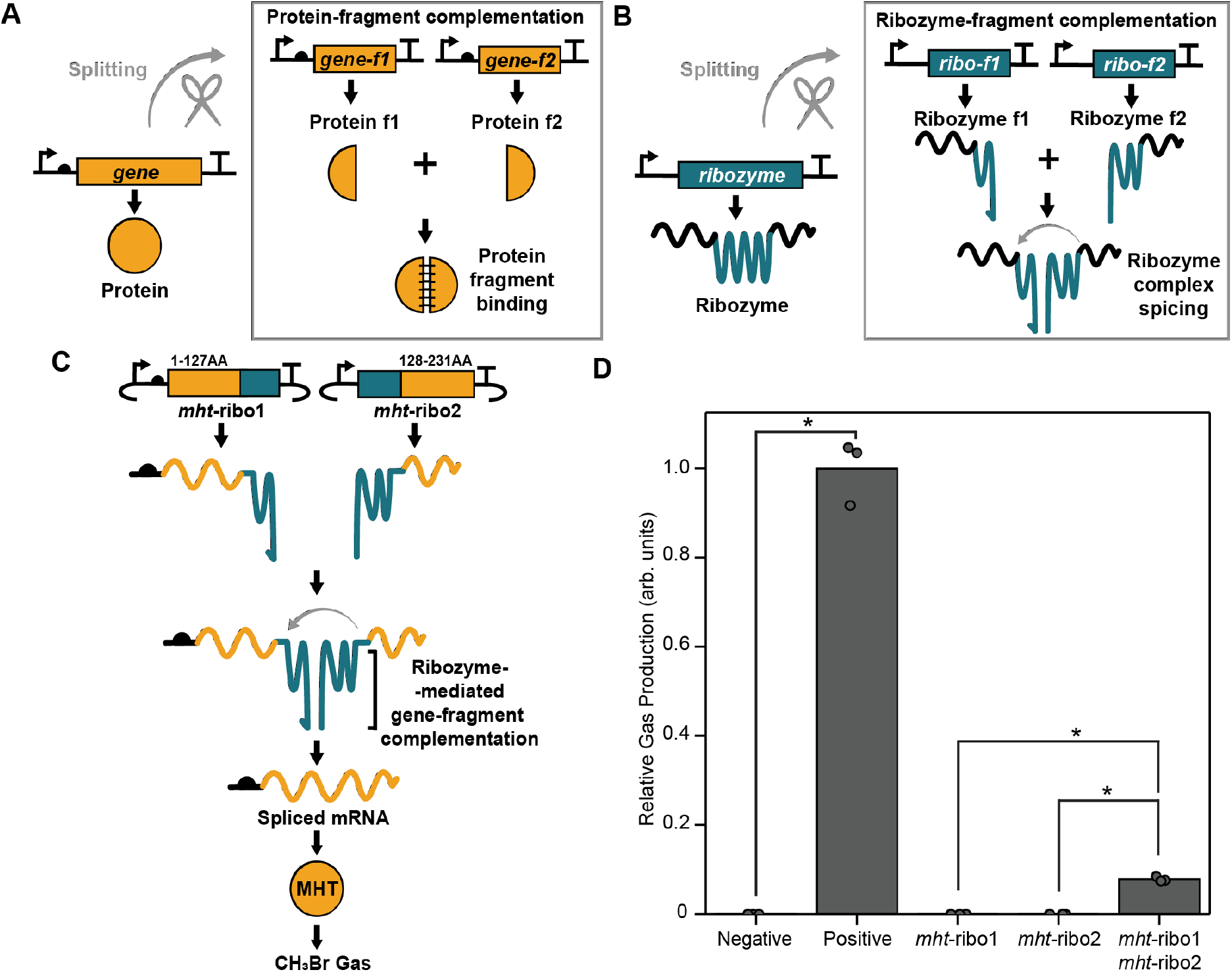
Combined protein and RNA control over reporter production. (**A**) Protein- and (**B**) ribozyme-fragment complementation can be combined to achieve (**C**) ribozyme-mediated gene-fragment complementation. This combined approach provides multi-layer control of an MHT reporter by splitting it at the protein and mRNA levels. (**D**) Relative gas production by *E. coli* MG1655 transformed with an empty plasmid (Negative), a plasmid that constitutively expresses MHT (Positive), and plasmids encoding each *mht*-ribozyme fragment (*mht*-ribo1 and *mht*-ribo2) alone or together. The relative gas production represents the ratio of CH_3_Br to CO_2_ scaled to the mean of the positive control. Biological replicates (n = 3) are shown as points. A one-way ANOVA of all data reveals gas production varies significantly between groups (F_6,14_ = 631, p < 3.3E-16). A Tukey’s HSD Test for multiple comparisons indicates that the signal from cells transcribing both fragments is significantly higher than cells transcribing the individual *mht*-ribozyme fragments (p-values = 0.047). Asterisks designate p-values < 0.05.

## Results and Discussion

### Controlling MHT activity with a split ribozyme

To achieve multi-level synthetic gene control over an MHT reporter, we split the gene encoding *Batis maritima* MHT at a site previously found to inactivate the enzyme^27^, and we fused the resulting *mht* fragments to *T. thermophila* ribozyme fragments that were found to be non-functional when expressed individually.^26^ By fusing RNA interaction sequences onto each *mht*-ribozyme fragment, the resulting pair of transcripts were designed to associate and enable ribozyme-mediated splicing, thereby attaching the *mht* gene fragments to create a transcript that translates full-length MHT (Figure 1C). To investigate if this design results in tight control over MHT translation and CH_3_Br production, the genes encoding the *mht*-ribozyme fragments (designated *mht-*ribo1 and *mht-*ribo2) were cloned onto separate DNA plasmids and transformed into *E. coli* MG1655 cells. Headspace gas analysis of cultures transformed with both plasmids revealed a CH_3_Br signal (Figures S1, 1D). In contrast, CH_3_Br was not detected with cells transformed with plasmids expressing the individual *mht-*ribozyme fragments. The signal from cells transcribing both *mht-*ribo1 and *mht-*ribo2 was ∼6% of that level observed with cells expressing the full-length MHT. In addition, the rate of indicator gas production remained linear for 2.5 hours, and the indicator gas production was proportional to cell counts (Figure S2). Taken together, these results show that ribozyme-mediated gene-fragment complementation yields stringent control over an indicator gas reporter.

To understand how temperature influences ribozyme-mediated gene-fragment complementation (Figure 2A), we evaluated performance across a range of temperatures. Since the temperature dependence of the indicator gas signal depends upon both ribozyme and MHT catalysis, we used a full-length MHT as a control (Figure 2B), which was constitutively expressed from the genome. Across all temperatures tested, 22 to 40°C, CH_3_Br was synthesized by *E. coli* MG1655 expressing the *mht*-ribozyme fragments and the native MHT (Figures S3). At each temperature, cells expressing the full-length MHT presented higher CH_3_Br signals than cells expressing the pair of *mht*-ribozyme fragments. To evaluate the effect of temperature on the split ribozyme activity, whole cell gas production was normalized to that observed at the highest temperature (Figure 2C). With this analysis, both designs presented similar relative activities across all three temperatures. This finding suggests that the ribozyme-mediated gene-fragment complementation is similar across the temperature range assayed.

**Figure 2.**
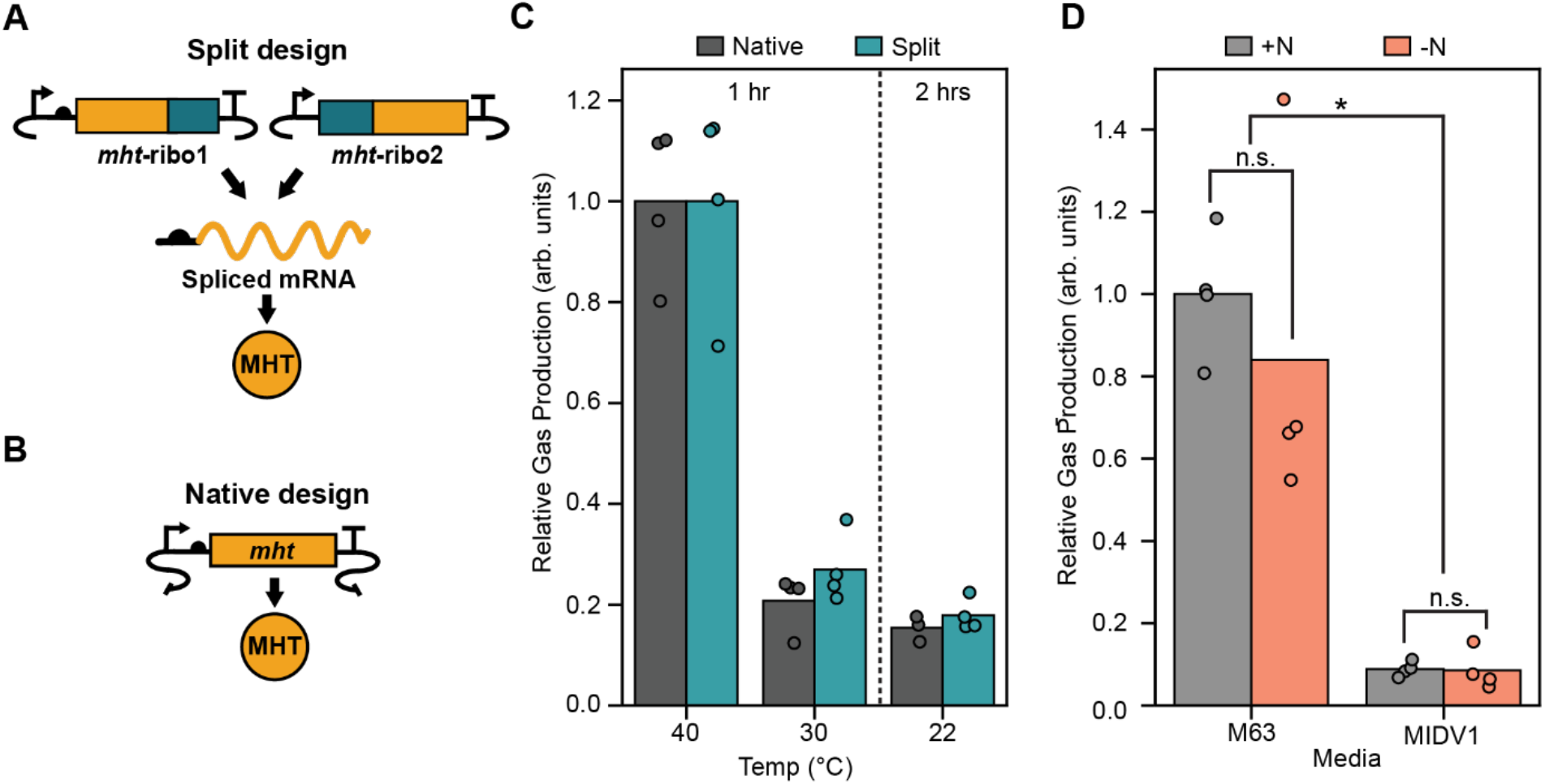
Effect of growth conditions on the indicator gas signal. The gas produced by cells expressing (**A**) both *mht*-ribozyme fragments and (**B**) full-length MHT were evaluated across (**C**) different temperatures. The relative gas signal represents the ratio of CH_3_Br to CO_2_ normalized to the average for each cell type at 40°C. Data represents 3 or more biological replicates. (**D**) Gas produced by cells expressing both *mht*-ribozyme fragments grown in M63 or MIDV1 containing (+N) or lacking (-N) nitrogen. Data is normalized to the highest value observed and represents four biological replicates. A one-way ANOVA showed gas production varies significantly between groups (F_3,13_ = 18.15, p < 9.33E-5). A Tukey’s HSD Test for multiple comparisons determined that gas production is not significantly impacted by nitrogen (p-values = 0.757 in M63 and 1.0 in MIDV1), and it found that gas production is significantly higher in M63 medium (p-values = 0.001 with N and 0.003 in the absence of N). The asterisk designates a p-value < 0.05.

To investigate whether ribozyme-mediated gene-fragment complementation can be used in growth medium that has soil-relevant osmolarity and nutrient conditions, we investigated the gas production of cells expressing the pair of *mht*-ribozyme fragments in M63 growth medium as well as a diluted version of this growth medium (MIDV1), which better mimics the osmolarity of a typical soil.^28^ Cells grown in M63 medium presented higher gas production than cells grown in MIDV1 medium (Figures S4, 2D). When gas production was evaluated in M63 and MIDV1 media lacking nitrogen, a similar signal was observed. Taken together, these results show that ribozyme-mediated gene-fragment complementation can be used in a minimal growth medium having an osmolarity and nitrogen depleted conditions that mirror soil conditions.

### Reporting on conjugation

We next sought to investigate if ribozyme-mediated gene-fragment complementation is efficient enough to report on conjugative gene transfer. To report on conjugation, *E. coli* MFDpir (donor) was transformed with a conjugative plasmid that transcribes *mht-*ribo1, and this strain was mixed with an *E. coli* MG1655 (recipient) that transcribes *mht-*ribo2 using a genetic circuit coded in the genome (Figure 3A). As a control, the latter strain was transformed with the conjugative plasmid to create a transconjugant. When present at the same titer as the recipient cells, the gas produced by this transconjugant represents the maximum potential gas signal if conjugation was 100% efficient and all recipient cells in a population received the plasmid (Figure S5). To assay conjugation, a series of cultures were incubated on rich agar medium supplemented with diaminopimelic acid (DAP) within sealed vials (2 mL), including (D) donor cells, (R) recipient cells, (D+R) donor and recipient cells, and (D+T) donor and transconjugant cells. DAP was included as it is required for donor cell growth since this strain lacks the *dapA* gene. Following incubation for two and five days (Figure 3B), CH_3_Br was not detected in vials containing either the donor or the recipient cells individually. In contrast, CH_3_Br was observed in vials containing both the donor and recipient cells. When this signal was normalized to the sample containing the donor and transconjugant cell mixture, which represents 100% conjugation efficiency, the signal corresponded to 0.29% ±0.25% (day 2) and 0.66% ±0.30% (day 5) of that observed with the transconjugant control. These findings show that ribozyme-mediated gene-fragment complementation reports on conjugation in *E. coli*, and it illustrates how multi-level synthetic gene control over an MHT reporter provides stringent control over gas synthesis in the absence of conjugation.

**Figure 3.**
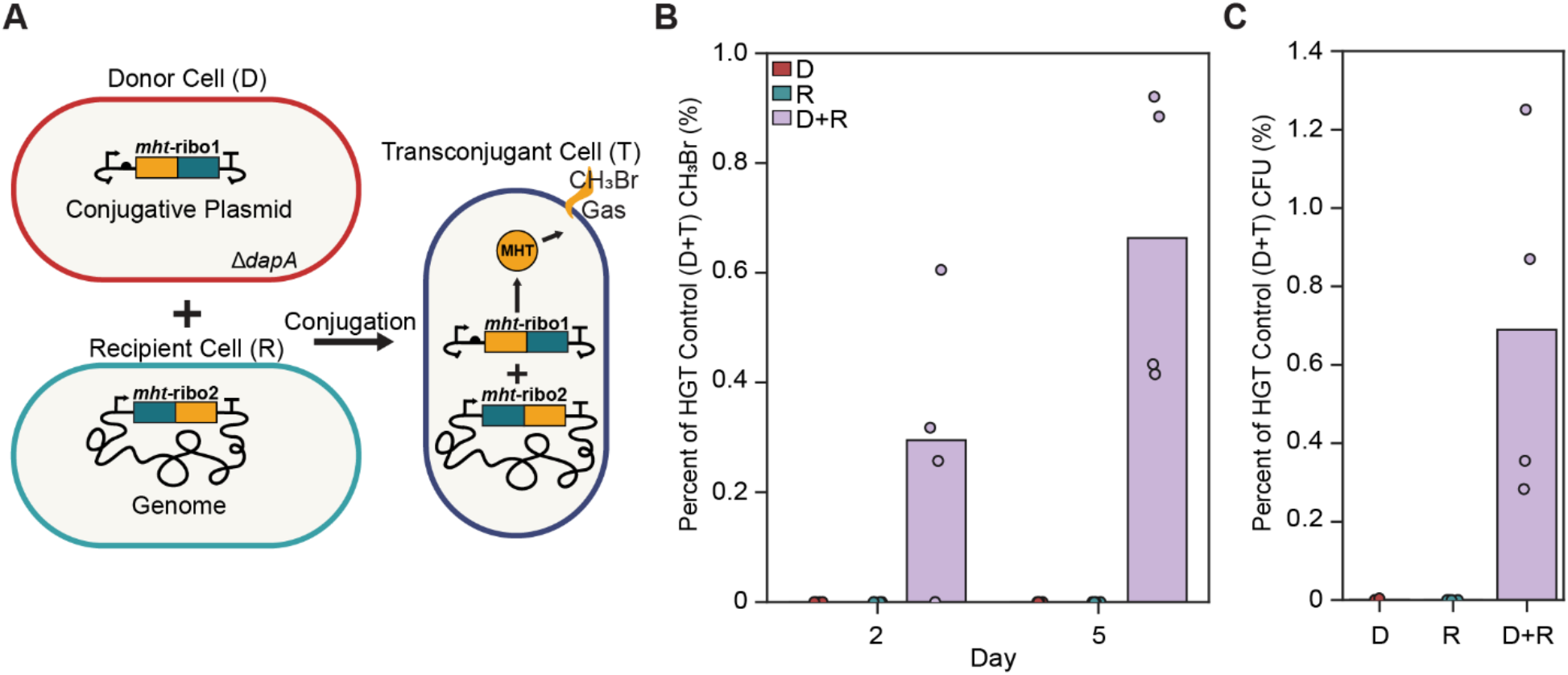
Comparing gas reporting of conjugation with antibiotic selection. (**A**) Strains used to perform conjugation. An *E. coli* donor (D) transformed with a conjugative plasmid encoding *mht*-ribo1 was mixed with an *E. coli* recipient (R) containing the *mht*-ribo2. Upon conjugation, transconjugant cells (T) are formed that synthesize CH_3_Br. (**B**) Whole cell gas production for D, R and D+R samples is scaled to the average signal from D+T samples. Data represents four biological replicates. A one-way ANOVA showed that CH_3_Br production varied significantly between groups on both day two (F_2,10_ = 5.64, p < 0.026) and five (F2,10 = 22.99, p < 0.0003). A Tukey’s HSD Test for multiple comparisons showed that the D+R signal was significantly higher than each control on days two and five (p-values = 0.04 and 0.001, respectively). (**C**) Transconjugant CFU measured on day five. Data from four biological replicates is scaled to the D+T sample average. A one-way ANOVA showed that CFUs varied significantly between groups (F_2,10_ = 9.13, p < 0.0068). A Tukey’s HSD Test for multiple comparisons found that CFUs from conjugation samples (D+R) were significantly higher than each control (p-values = 0.012).

To investigate how the CH_3_Br signal arising from mixing the donor and recipient cells relates to the conjugation rate, we quantified conjugation by selecting for transconjugants on agar plates using antibiotics, a commonly used approach to monitor conjugation.^29,30^ To allow for a direct comparison, we resuspended cells from the headspace gas measurements (day five) in liquid medium and plated them on agar plates containing a pair of antibiotics (kanamycin and chloramphenicol) that select specifically for the recipient cells [Cm^R^] that acquired the conjugative plasmid [Kan^R^], which represent transconjugant cells. Figure 3C shows that the number of colony forming units (CFU) observed with the donor and recipient (D+R) mixture was 0.69% ±0.56%. This value is similar to that calculated using the headspace gas analysis (0.66% ±0.30%). A comparison of the individual biological replicates revealed a correlation between the number of transconjugants observed using antibiotic selection and gas production (Figure S6). Taken together, these findings show that ribozyme-mediated gene-fragment complementation yields conjugation efficiency information that mirrors traditional selection approaches.

### Detecting conjugation in soil

We next sought to investigate if ribozyme-mediated gene-fragment complementation can report on gene transfer when conjugation is performed within a soil, which can present a background CH_3_Br signal.^24^ Conjugation measurements were performed in a previously described soil (Figure 4A) that had been hydrated to ∼20% water holding capacity (WHC).^23^ Prior to performing conjugation experiments, we evaluated donor cell viability upon adding different DAP concentrations (Figure S7). To remain viable in soil, donor cells required a higher DAP concentration than in liquid medium, indicating that soil decreases DAP bioavailability.

**Figure 4.**
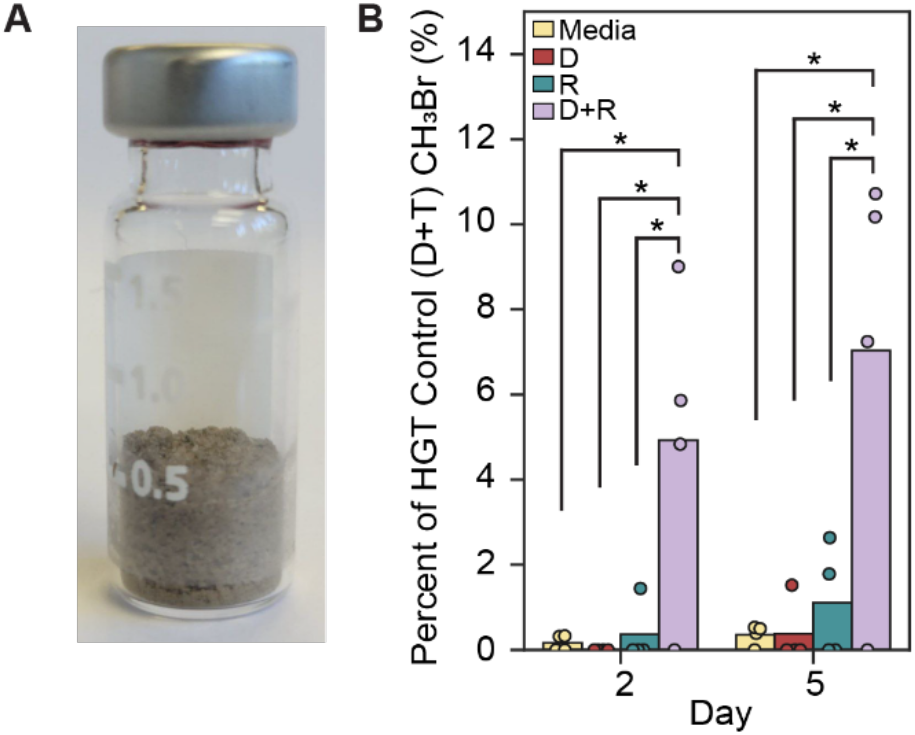
Conjugation-induced gas production in soil. (**A**) Vials containing B Horizon soil were hydrated to 20% WHC using growth medium (Media) or medium containing donor (D), recipient (R), or donor and recipient cells (D+R). (**B**) Signals were scaled to the average from donor and transconjugant mixtures (D+T). Data represents four biological replicates. A one-way ANOVA showed that gas production varied significantly between groups on both day 2 (F_3,13_ = 6.259, p < 0.008) and day 5 (F_3,13_ = 6.249, p < 0.008). A Tukey’s HSD Test for multiple comparisons found that the D+R signal was significantly higher than other samples on day two (p-values = 0.018 for media, 0.015 for D, and 0.024 for R) and five (p-values = 0.015 for media, 0.015 for D, and 0.031 for R).

To study conjugation in soil, *E. coli* MFDpir (donor) transformed with the conjugative plasmid that transcribes *mht-*ribo1 was mixed with *E. coli* MG1655 (recipient) that transcribes *mht-*ribo2 from a plasmid. A two-plasmid system was used to enhance the copy number of the latter transcript, which was critical for observing a robust gas signal from transconjugants in soil (Figure S8). When the soil was hydrated in the absence of donor or recipient cells, a background CH_3_Br signal was observed (Figures 4B, S9), similar to prior studies^23^. Soil containing the donor or recipient strains presented a gas signal with a similar magnitude. In contrast, soil containing both the donor and recipient cells presented significantly higher CH_3_Br. When this CH_3_Br signal was compared to the transconjugant control, the CH_3_Br levels were 4.9 ±4.2% and 7.0 ±6.8% of that observed with the transconjugant following two- and five-day incubations, respectively. These findings show that ribozyme-mediated MHT-fragment complementation can be used to report on conjugation within soils, which present background CH_3_Br.

To remove the requirement for DAP in conjugation studies, whose bioavailable levels were influenced by soil, a conjugative plasmid was created that transcribes *mht*-ribo1 and expresses DapA. To investigate if *E. coli* MFDpir (donor) transformed with this plasmid presents enhanced fitness compared with cells harboring the original conjugative vector, growth was evaluated in liquid medium containing or lacking DAP (Figure 5A). With this modified donor strain, growth was significantly enhanced in the absence of DAP. We also evaluated whether the conjugative plasmid containing the *dapA* gene supports gas production in transconjugant cells (Figure 5B, S10). CH_3_Br was observed with cells containing the conjugative plasmid with the *dapA* gene, although it was decreased ∼50% compared to the original plasmid. This finding suggested that this plasmid would be sufficient to enable the detection of conjugation without requiring DAP addition to soil.

**Figure 5.**
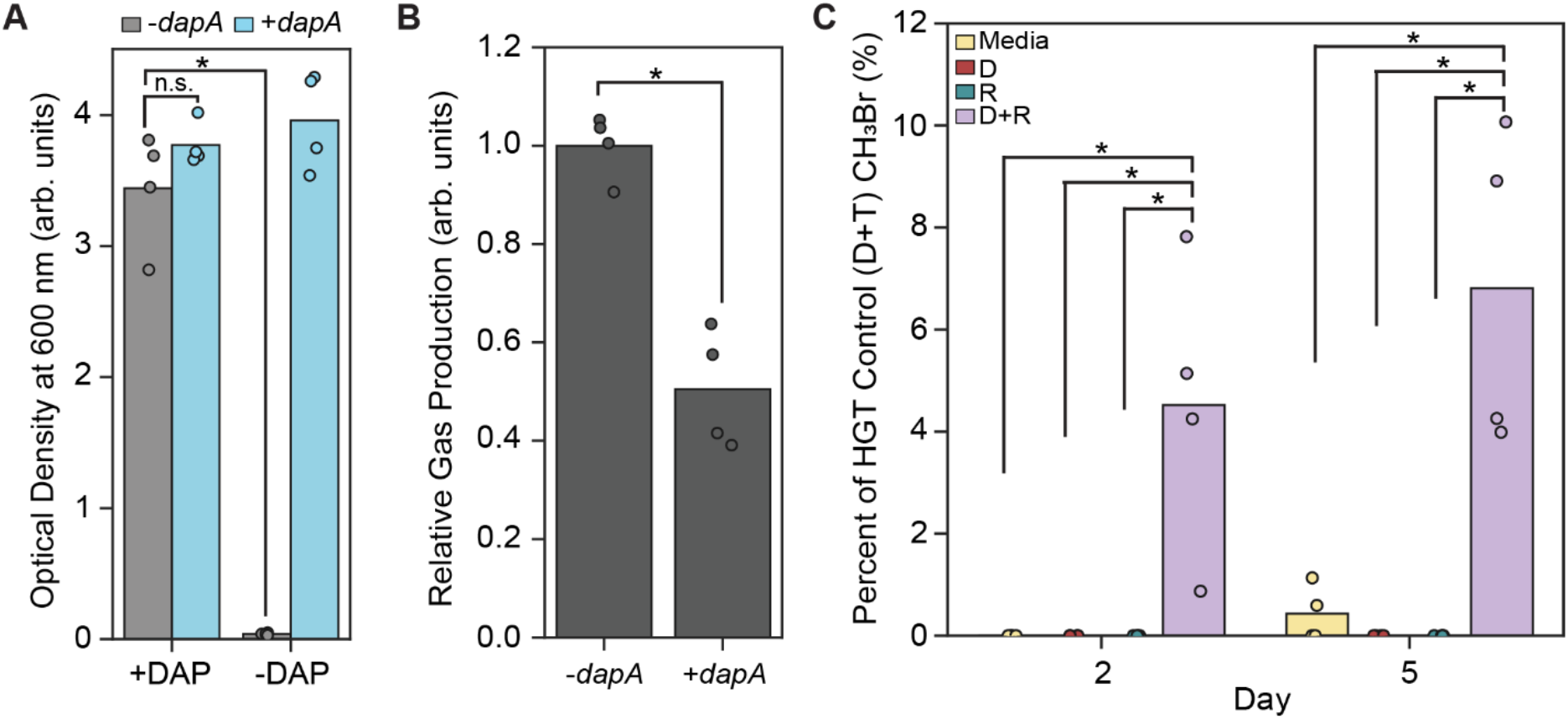
Monitoring conjugation in soil in the absence of DAP. (**A**) Effect of DAP on *E. coli* MFDpir transformed with a plasmid containing or lacking *dapA*. Following overnight growth at 37°C, optical density at 600 nm (OD_600_) was measured. A one-way ANOVA showed OD_600_ varied significantly between groups (F_3,13_ = 151.9, p < 8.3E-10). A Tukey’s HSD Test for multiple comparisons revealed that cells lacking *dapA* present an OD_600_ that is significantly decreased by DAP (p-value < 0.001). In the absence of DAP, the +*dapA* strain grew significantly better than the -*dapA* strain (p-value < 0.001). (**B**) Gas production by transconjugants containing or lacking *dapA* in liquid medium. The gas production from the +*dapA* strain was significantly higher than the *-dapA* strain (t_6_ = 7.217, p < 0.001). (**C**) Headspace gas in soils lacking DAP that were hydrated with growth medium (Media) and medium containing donor (D), recipient (R), or both donor and recipient (D+R) cells. Data are reported as a percentage of the average signal observed with soil containing a mixture of D+T. A one-way ANOVA showed that CH_3_Br varied significantly between groups on both day two (F_3,13_ = 9.94, p < 0.0014) and five (F_3,13_ = 17.58, p < 0.0001). A Tukey’s HSD Test revealed that CH_3_Br production from conjugation samples was significantly higher than each control sample for both day two (p-value = 0.004 for media, donor, and recipient) and day five (p-value = 0.001 for media, donor, and recipient). Data represents four biological replicates, and asterisks indicate p-values < 0.05.

To determine if the modified reporter system could report on HGT in soil lacking DAP, conjugation was repeated in soil hydrated to 20% WHC. Soil containing donor or recipient cells all presented low CH_3_Br levels (Figures 5C, S11), like that observed with soil hydrated using medium alone. In contrast, the samples containing both donor and recipient cells produced significantly more CH_3_Br, with levels that were ≥16-fold higher than the controls following a 5-day incubation. A comparison of the CH_3_Br levels from the donor and recipient sample to the transconjugant control revealed a signal that was 4.5 ±3.0% and 6.8 ±4.2% of the positive control on days two and five, respectively. These results show how ribozyme-mediated gene-fragment complementation can report on conjugation in soil without DAP supplementation.

### Implications

Our results illustrate how multi-level synthetic gene control can be achieved over indicator gas production using ribozyme-mediated gene-fragment complementation. This approach enables exquisite control over MHT translation arising from gene transfer, thereby yielding a gas reporter of conjugation that is compatible with applications within hard-to-image soils. This finding can be contrasted with prior applications of gas reporters in soil, which found it challenging to repress the output signal in the absence of gene transfer.^25^ The gas reporting approach described herein has advantages over other reporters used in soils, such as visual and ice nucleation reporters.^31–33^ First, it produces a signal that can be measured in the headspace of bulk soil samples without requiring microbial extraction, which is arduous. This characteristic makes this reporting approach compatible with studies within diverse environmental materials, ranging from soils and sediments to wastewater and sludge, as it can be used in safe containers commonly used by environmental scientists to monitor the temporal dynamics of greenhouse gas production.^34,35^ Second, MHT functions as a reporter under both aerobic and anaerobic conditions.^19^ This can be contrasted with more traditional visual reporters like GFP, which require oxygen to mature into a fluorescent protein.^32^

Currently, our understanding of soil controls on gene transfer remains limited, although it is recognized that the biological and physicochemical characteristics of soil will influence rates.^36^ In future studies, ribozyme-mediated gene-fragment complementation should allow for measurement of rare HGT events in real soils across day, week, and month long incubations. Since this tool has been shown to have tight control over gas production across a wide range of nutrient conditions and temperatures, we expect that this HGT reporter will aid in studying how soil matrix parameters impact HGT using defined pairs of microbes. For example, it can be used to study how anthropogenic chemicals, like antibiotics^37^, affect gene transfer dynamics *in situ*. In addition, it will be useful for studying how physical properties of the soil, such as matric potential, influence gene transfer. By establishing how different soil properties control gene transfer, HGT models can be calibrated for real environmental settings^38^, which will be important for understanding how climate change may alter gene transfer in soil environments. Furthermore, this tool will allow for evaluation of biocontainment approaches developed for synthetic biology^39^, whose performance remains uncertain in these settings. Such knowledge will allow for a more wholistic understanding of HGT in situ and the risks of deploying genetic-engineered technologies in soils.^40^

## Acknowledgements

This research was supported by the United States Department of Agriculture Biotechnology Risk Assessment grant 2021-33522-35356 (to JC and JJS) and National Science Foundation grant 1828869 (to JJS). The authors thank Ella Whitehead and Dr. Li Chieh Lu for technical support and Dr. Andrea Garza for constructive discussions. Additional thanks to the Masiello lab for support with GC-MS measurements.

## Methods

### Plasmids and Strains

Plasmids are listed in Table S1, strains in Table S2, and sequences in Table S3. Plasmids were constructed using a combination of Gibson, Golden Gate, and inverse PCR assembly.^41–43^ Plasmid construction was performed using NEB Turbo *E. coli*, while reporter characterization used strains derived from *E. coli* MG1655. Genome integration of *mht*-ribo2 and a chloramphenicol resistance (CmR) cassette into *E. coli* MG1655 was performed at the *lacZ* loci.^44^ All DNA modifications were confirmed by Sanger sequencing. The *E. coli* MFDpir strain has genomically integrated RP4 conjugation machinery allowing for plasmid donation, and it requires DAP supplementation for growth.^45^

### Growth Medium

Lysogeny Broth (LB) was used for all molecular biology. M63 and MIDV1 were prepared as in Table S4 within 3 days of use by combining sterile ingredients and supplementing with 100 mM NaBr. To create nitrogen-free media, M63 and MIDV1 were prepared without casamino acids and using an M63 5x salt mixture lacking ammonium sulfate. For GC-MS studies in nutrient-rich conditions, LB was used that contains NaBr (100 mM) rather NaCl, designated LB-Br (Table S4). Medium with DAP was prepared by adding sterile DAP (60 mM) to the desired concentrations (Table S5).

### Transformation

Cells were transformed using chemical or electrocompetent cells. Chemically-competent *E. coli* NEB Turbo and MG1655 (50 µL) were mixed with plasmid (up to 10 µL), incubated on ice for 20 to 40 minutes, heat shocked at 42°C for ∼1 minute, iced for 2 minutes, mixed with LB (50 µL), and incubated for one hour at 37°C. After spreading on LB-agar with antibiotics, cells were incubated overnight at 37°C. *E. coli* MFDpir were electroporated with the EC1 setting on a Bio-Rad MicroPulser using a 0.1 cm gap electroporation cuvette and mixing the cells with LB (1 mL). After incubating ∼1 hour at 37°C, cells were spread on LB-agar containing antibiotics. Following overnight growth, single colonies were used to inoculate cultures.

### Conjugation measurements

Conjugation was performed by growing cultures overnight at 37°C in LB containing kanamycin (100 µg/mL, donor strain), chloramphenicol (34 µg/mL, genome-encoded recipient strain), or spectinomycin (50 µg/mL, plasmid-encoded recipient strain). Cultures contained 0.3 mM DAP when needed for growth. Then, cultures were washed ≥3 times using LB. To perform conjugation on agar, LB-Br agar medium was prepared as in Table S4, autoclaved, and supplemented with DAP (0.3 mM). Before solidifying, hot medium (200 µL) was added to vials (2 mL), vials were capped, and vials were stored at 4°C until use. To initiate conjugation, donor and recipients were mixed at a 1:1 ratio by volume and spotted onto LB-agar containing DAP (0.3 mM) for 2 and 5 days at 37°C; vials were immediately capped. Following methyl bromide analysis of conjugation (day five, Figure 3B), cells were resuspended in LB medium, serially diluted, and spread onto LB-agar medium containing chloramphenicol (34 µg/mL) and kanamycin (100 µg/mL). After overnight incubation at 37°C, colonies were counted and used to calculate CFU. To perform conjugation in soil, all steps were identical except cells were not removed from media post-gas measurements for CFU counting. In cases where a plasmid-encoded recipient cell was used for conjugation, selection plates for transconjugants contained kanamycin (100 µg/mL) and spectinomycin (50 µg/mL). HGT percent was calculated by taking the average CH_3_Br production from donor mixed with recipient (D+R samples) and dividing by the average CH_3_Br production for that day from the positive control (donor mixed with transconjugant, D+T samples) (Table S6).

### GC-MS Sample Preparation

Figure S12 shows an overview of the protocol, while Table S5 describes the individual parameters that varied by experiment. Samples were prepared by picking individual colonies from plates into the pre-culture media containing antibiotics and DAP if needed for strain growth. Cultures were then grown overnight in a shaking incubator (225 rpm). Cultures (500 µL) were pelleted by centrifugation (1.5 min at 6010 RCF, 4°C), resuspended in 1000 mL of incubation medium, and this process repeated three times. The samples were diluted to a set optical density at 600 nm (OD_600_) and placed on ice. Vials containing samples were then capped using a protocol that avoided variation in incubation time. For sample preparation in vials, cells were added to the GC-MS vials that were either empty or contained solid medium. When a mixture of two cultures was used in a single sample, *e*.*g*., donor and recipient cells, cultures were mixed prior to addition to GC-MS vials. Following sample addition, vials were crimped with a cap and incubated without shaking prior to GC-MS analysis.

For experiments using different temperatures, cells were diluted to an OD_600_ of 0.04 and grown until they reached 3 doublings (∼0.32). Due to the differences in growth rates across temperatures, the sample at 22°C was incubated for 2 hours rather than 1 hour to reach a similar post-measurement OD_600_. For experiments exploring different growth media, pre-cultures were prepared in the nitrogen-containing versions of each media type. This approach minimized the metabolic shift when cells were transferred to the test media since MIDV1+N pre-cultures were used to prepare both MIDV1+N and MIDV1-N cultures. M63 and LB-Br sample preparation were similar. All steps post-washing followed the general GC-MS sample preparation protocol.

### GC-MS Analysis

The GC-MS used (Table S5) included: (i) an Agilent 7820A (GC) with an Agilent 5977E (MS) and an Agilent 7693A autosampler fitted with a 100 μL gastight syringe was used to inject headspace gas (50 μL) into either a PoraPLOT Q capillary column (25 m, 0.25 mm ID, 8 μm film; Agilent) or a DB-VRX capillary column (20 m, 0.18 mm ID, 1 μm film; Agilent); (ii) an Agilent 7890B (GC) with an Agilent 5977B (MS) and an Agilent 7693A autosampler fitted with a 100 μL gastight syringe was used to inject headspace gas (50 μL) into a DB-VRX capillary column (20 m, 0.18 mm ID, 1 μm film; Agilent), and (iii) an Agilent 8890 (GC) with an Agilent 5977B (MS) and an Agilent 7693A autosampler fitted with a 100 μL gastight syringe was used to inject headspace gas (50 μL) into a DB-VRX capillary column (20 m, 0.18 mm ID, 1 μm film; Agilent). Data analysis was performed using Agilent MassHunter WorkStation Quantitative Analysis, which determined the peak area for CH_3_Br using only CH_3_^79^Br, which contains the major isotope. To check for compound identity, the ratio between CH_3_^79^Br and CH_3_^81^Br was evaluated, especially at lower peak areas. All vials (2 mL) and caps were from Phenomenex.

### Soil Preparation

All soil was from previously-characterized B Horizon soil collected in 2020 from Nature’s way in Conroe, Texas which had been baked, sieved, and stored at room temperature immediately following collection.^23^ Prior to experiments, soil was twice autoclaved. GC-MS vials were prepared by weighing out one gram of soil, adding it to the vial, and crimping. Vials were uncapped prior to sample addition.

