## Supporting Information for "Ribozyme-mediated gene-fragment complementation for non-destructive reporting of DNA transfer within soil"

### Table of Contents


| Supplemental Figure | Pages |
|---|---|
| Figure S1. Effect of split ribozyme on whole cell gas production | 3 |
| Figure S2. Kinetics of whole cell gas production | 4 |
| Figure S3. Effect of temperature on whole cell gas production | 5 |
| Figure S4. Effect of growth medium on whole cell gas production | 6 |
| Figure S5. Conjugation experiment sample types | 7 |
| Figure S6. Comparing antibiotic selection and gas reporting of conjugation | 8 |
| Figure S7. Effect of DAP concentration on cell growth | 9 |
| Figure S8. Genome and plasmid encoded reporters vary in signal | 10 |
| Figure S9. $CH_3Br$ production in soils containing DAP | 11 |
| Figure S10. Effect of the *dapA* gene on whole cell gas production | 12 |
| Figure S11. $CH_3Br$ production in soils lacking DAP | 13 |
| Figure S12. Overview of GC-MS sample preparation | 14 |

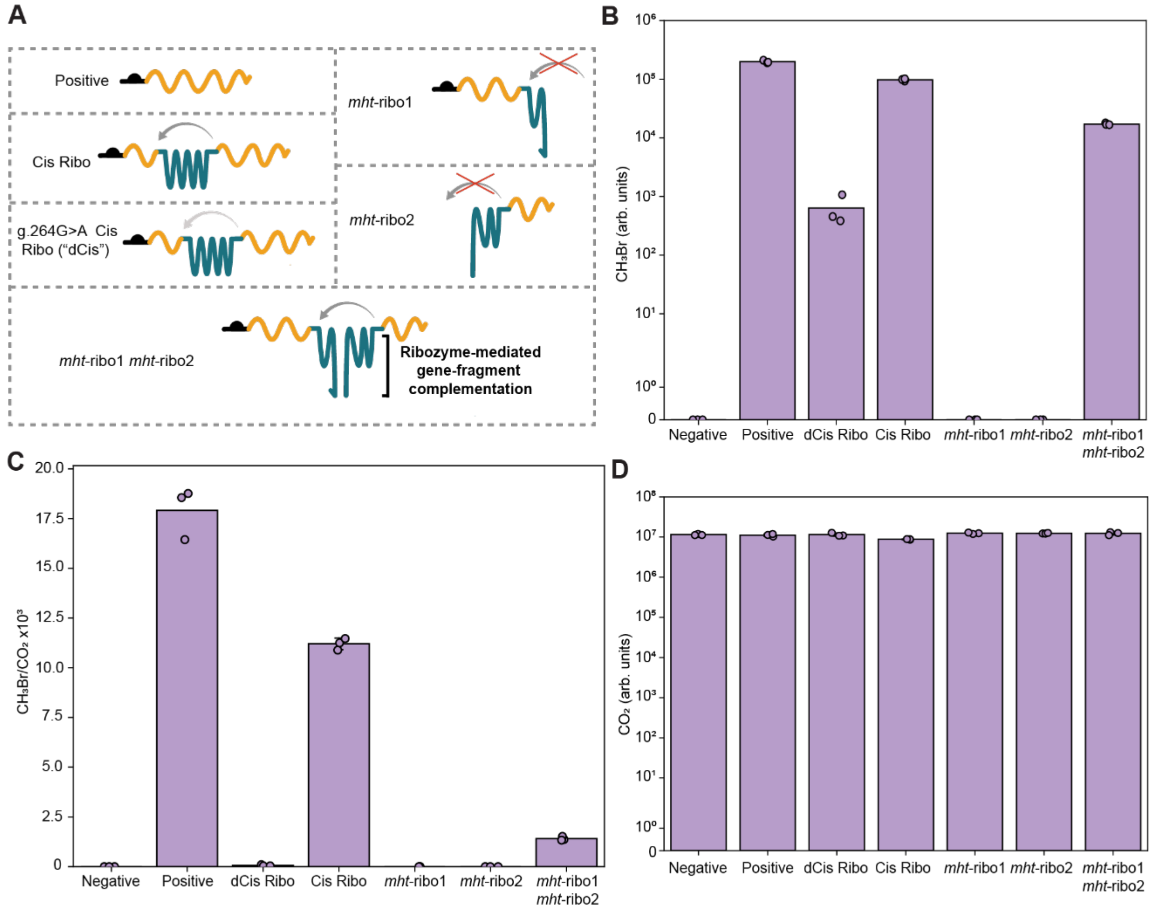


**Figure S1. Effect of split ribozyme on whole cell gas production.** (**A**) RNA designs tested included native *mht* (Positive), a self-splicing ribozyme-MHT (Cis Ribo), each ribozyme-*mht* fragment alone (*mht*-ribo1 or *mht*-ribo2), each ribozyme-*mht* fragment together (*mht*-ribo1 and *mht*-ribo2), or a self-splicing ribozyme-MHT with an inactivating point mutation in the ribozyme (g.264G>A, dCis Ribo). For *E. coli* MG1655 transcribing each design, we measured (**B**) $CH_3Br$ and (**C**) $CO_2$. These values were used to calculate (**D**) the ratio of the $CH_3Br/CO_2$ signals. Data represents three biological replicates for each measurement.

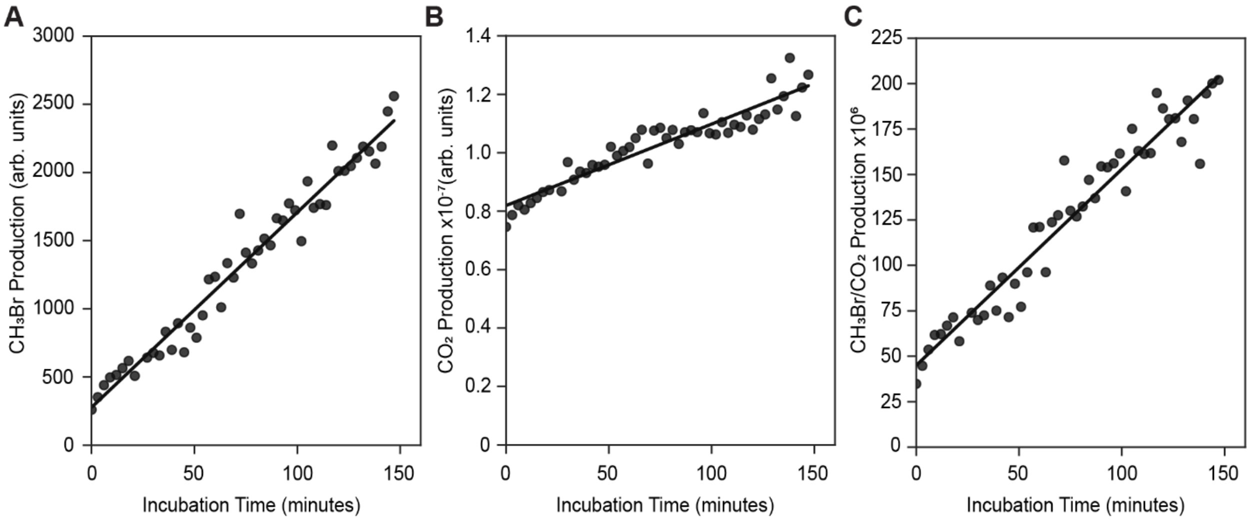


**Figure S2. Kinetics of whole cell gas production at room temperature.** (**A**) $CH_3Br$, (**B**) $CO_2$, and (**C**) the ratio of $CH_3Br/CO_2$ as a function of time for a single culture of *E. coli* MG1655 transcribing *mht*-ribo1 and *mht*-ribo2. Linear fits yield $R^2$ values of 0.96 ($CH_3Br$), 0.88 ($CO_2$), and 0.93 ($CH_3Br/CO_2$).

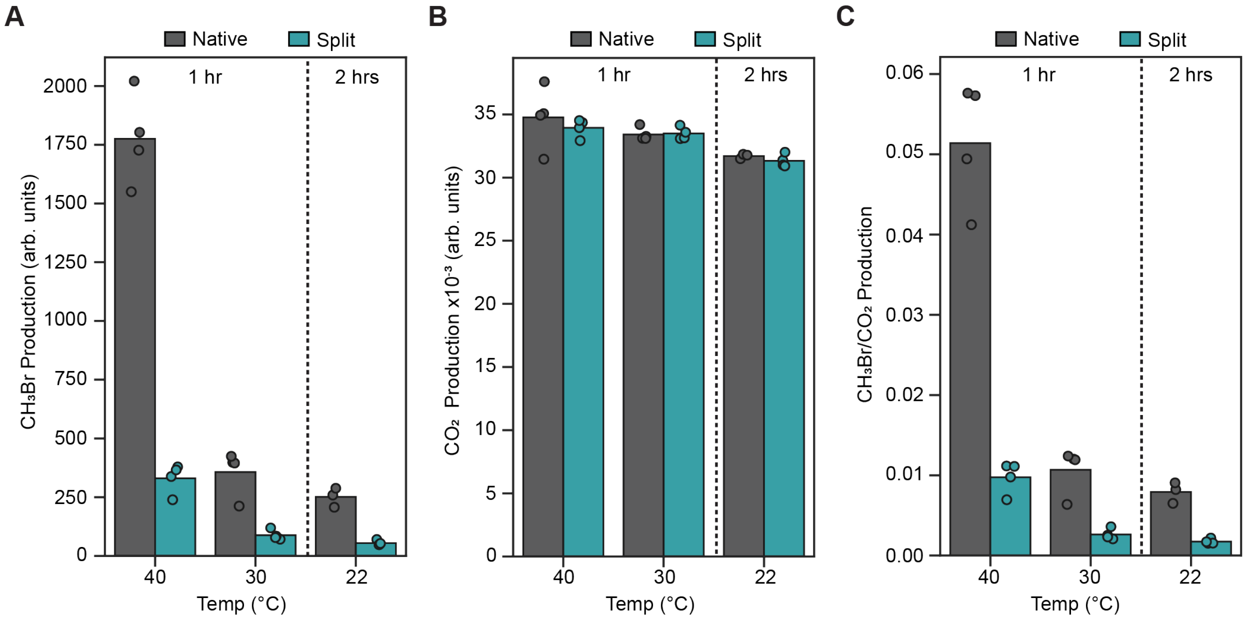


**Figure S3. Effect of temperature on whole cell gas production.** (**A**) $CH_3Br$, (**B**) $CO_2$, and (**C**) the ratio of $CH_3Br/CO_2$ produced by *E. coli* transcribing *mht*-ribo1 and *mht*-ribo2 (Split) or the full-length MHT gene (Native). Samples at 30 and 40°C were measured after a 1-hour incubation while samples at 22°C were analyzed after a 2-hour incubation. Data represents ≥3 biological replicates shown as points.

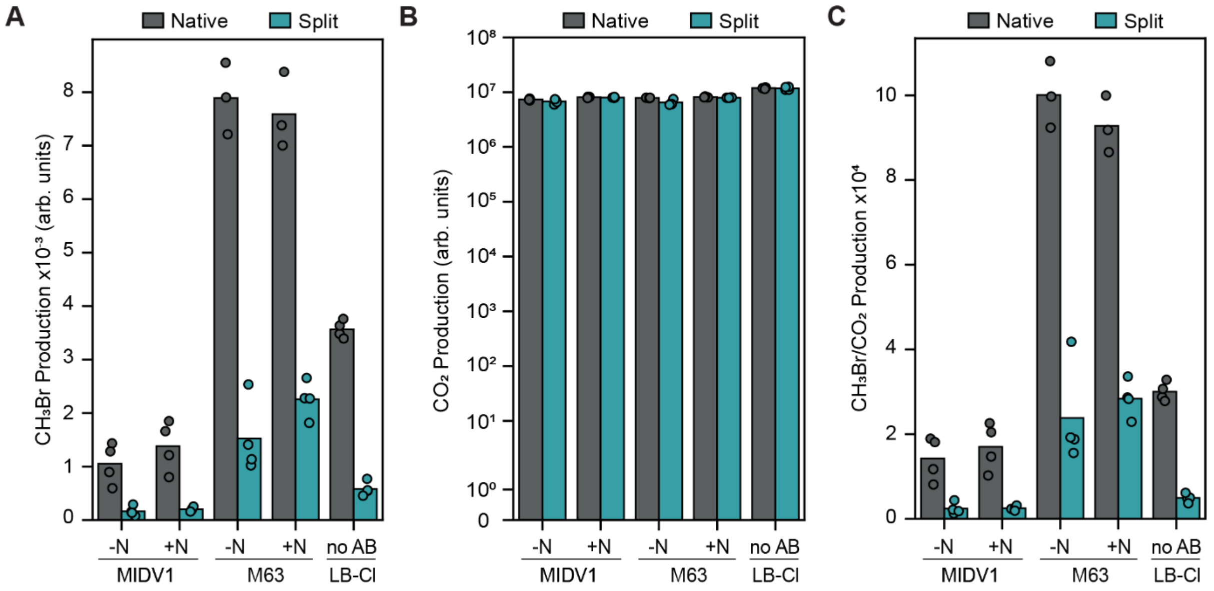


**Figure S4. Effect of growth medium on whole cell gas production.** (**A**) $CH_3Br$, (**B**) $CO_2$, and (**C**) the ratio of $CH_3Br/CO_2$ produced by *E. coli* transcribing *mht*-ribo1 and *mht*-ribo2 (Split) or the full-length MHT gene (Native). Three growth media were evaluated including M63, MIDV1, and LB-Br. With M63 and MIDV1, culture conditions contained (+N) or lacked (-N) nitrogen. The minimal media contained antibiotics while no antibiotic was used in LB-Br. Prior to gas measurements, cultures were diluted to an optical density at 600 nm ($OD_{600}$) of 0.5, capped, and incubated for 1 hour at 22°C prior to headspace gas analysis. Data represents four biological replicates shown as points.

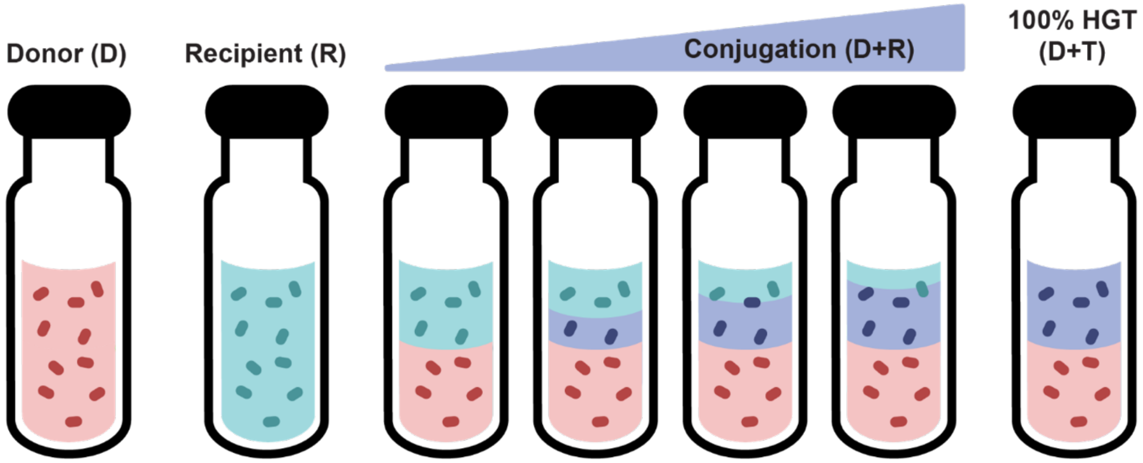


**Figure S5. Conjugation experiment sample types.** Vials contained either donor (D) alone, recipient (R) alone, a donor and recipient mixture (D+R), or a donor and transconjugant mixture (D+T). Conjugation was measured using a 1:1 ratio of D+R cells. To establish a frame of reference for T cells (purple), a control was generated with a 1:1 D+T mixture to mimic the gas signal if all R cells were converted to T cells.

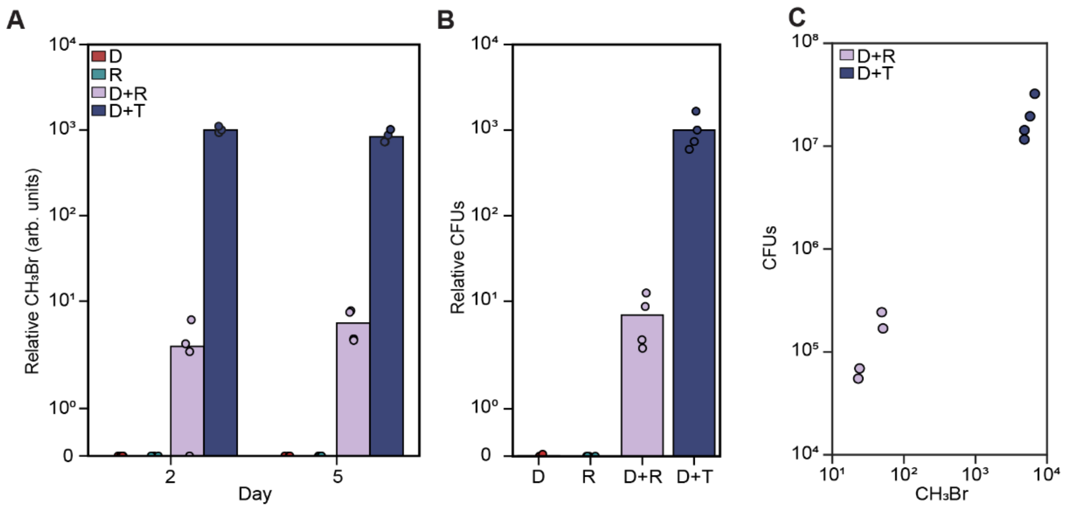


**Figure S6. Comparing antibiotic selection and gas reporting of conjugation.** (**A**) Gas produced by cells in capped vials where cells were spotted onto agar pads, including donor and recipient cells at 1:1 ratio (D+R), donor and transconjugant cells at a 1:1 ratio (D+T), donor (D) alone, and recipient (R) alone. (**B**) Transconjugants quantified by colony forming units (CFU) obtained when cell mixtures from panel A (day 5) were plated on LB-agar containing antibiotics. Relative indicator gas and CFU were calculated by scaling data to the mean of the highest D+T signal. (**C**) Comparison of relative indicator gas and CFU on day 5 for individual replicates. Data represents four biological replicates for each sample type shown as points.

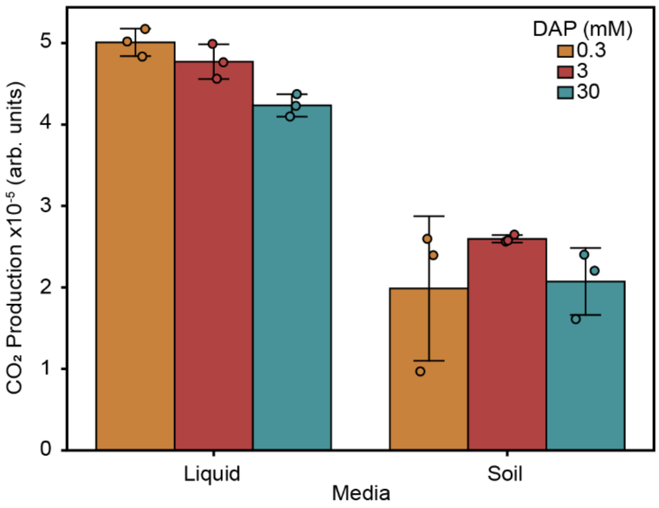


**Figure S7. Effect of DAP concentration on cell growth.** Effect of DAP concentration on the respiration of *E. coli* MFDpir transcribing *mht*-ribo1. Data represents three biological replicates shown as points and error bars show standard deviation.

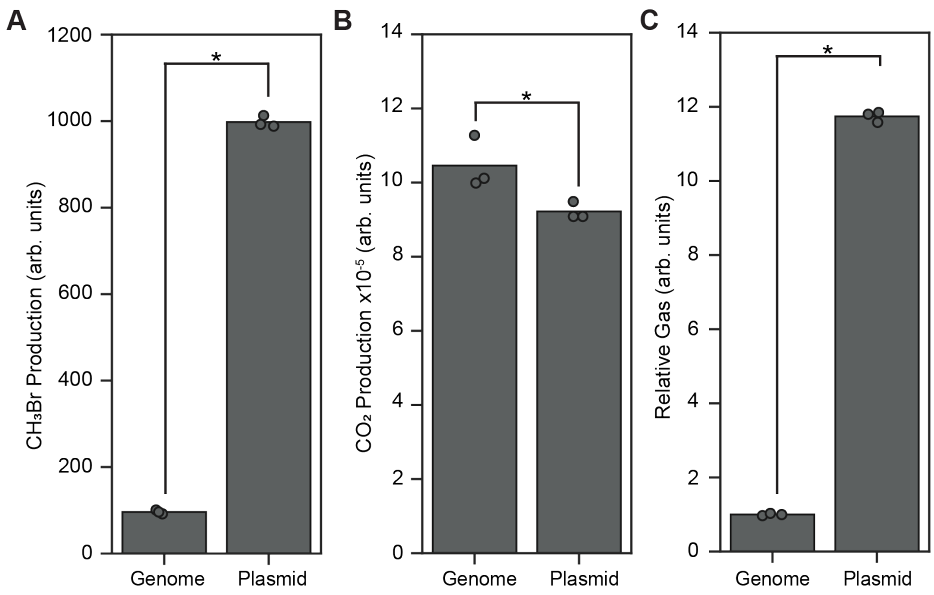


**Figure S8. Genome and plasmid encoded reporters vary in signal.** (**A**) $CH_3Br$, (**B**) $CO_2$, and (**C**) the ratio of $CH_3Br/CO_2$ (Relative gas) produced by *E. coli* that transcribes *mht*-ribo1 from a plasmid and mht-ribo2 from either the genome (Genome) or a plasmid (Plasmid). Relative gas production is reported relative to the average for the Genome sample. Data represents four biological replicates for each sample type shown as points. Vials inoculated with cells at an $OD_{600}$ of 0.5 in LB-Br were incubated at 37°C for 6 hours prior to headspace gas analysis. Asterisks designate p-values < 0.05.

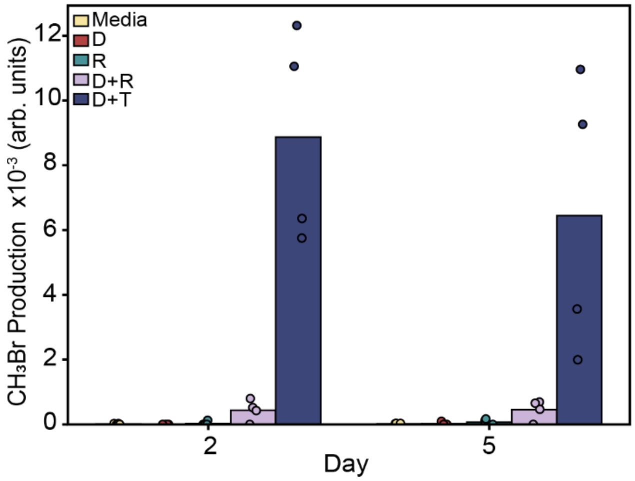


**Figure S9. $CH_3Br$ production in soils containing DAP.** Gas in the headspace of autoclaved B horizon soil that was hydrated with medium containing donor (D), recipient (R), donor and recipient at 1:1 ratio (D+R), or donor and transconjugant at a 1:1 ratio (D+T). Data represents four biological replicates shown as points.

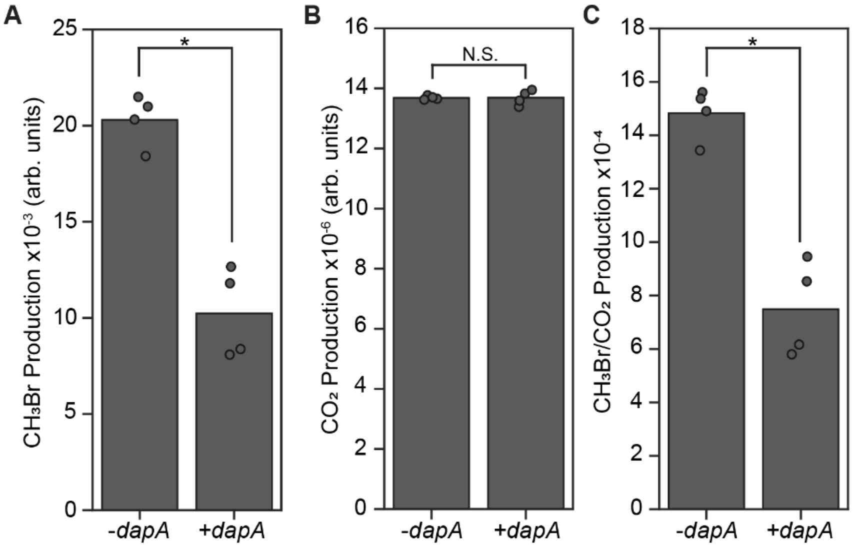


**Figure S10. Effect of the *dapA* gene on whole cell gas production.** (**A**) $CH_3Br$, (**B**) $CO_2$, and (**C**) the ratio of $CH_3Br/CO_2$ produced by *E. coli* MFDpir transcribing *mht*-ribo1 and *mht*-ribo2. Cells additionally contained (+*dapA*) or lacked (-*dapA*) a module for expressing DapA. Data represents four biological replicates shown as points. Asterisk designates p-values < 0.05. N.S., not significant.

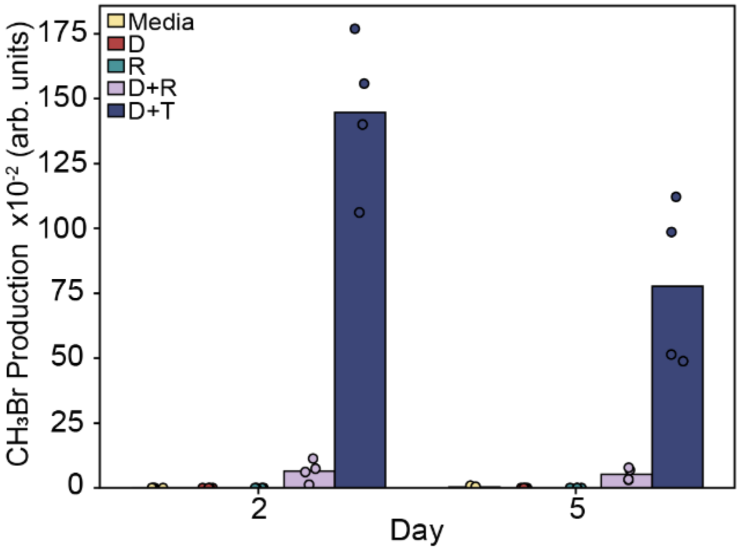


**Figure S11. $CH_3Br$ production in soils lacking DAP.** Gas in the headspace of autoclaved B horizon soil that had been hydrated with medium containing donor cells (D), recipient cells (R), donor and recipient cells at 1:1 ratio (D+R), or donor and transconjugant cells at a 1:1 ratio (D+T). Data represents four biological replicates shown as points.

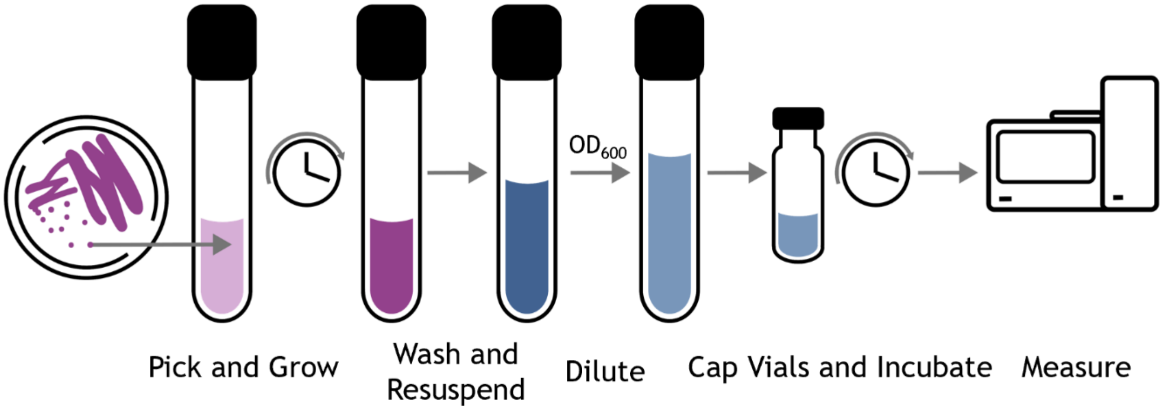


**Figure S12. Overview of GC-MS sample preparation.** Colonies from plates were used to inoculate a pre-culture (pink). After reaching stationary phase (purple), cells were washed and resuspended in the experimental medium (dark blue). This process was designed to remove antibiotics and chlorine from the pre-culture medium. The $OD_{600}$ was measured and used to dilute cultures to a desired density (light blue) within 2 mL vials. After capping, vials were incubated at the desired temperature prior to headspace gas analysis using GC-MS.
